# Nociceptor neuroimmune interactomes reveal cell type- and injury-specific inflammatory pain pathways

**DOI:** 10.1101/2023.02.01.526526

**Authors:** Aakanksha Jain, Benjamin M. Gyori, Sara Hakim, Samuel Bunga, Daniel G Taub, Mari Carmen Ruiz-Cantero, Candace Tong-Li, Nicholas Andrews, Peter K Sorger, Clifford J Woolf

## Abstract

Inflammatory pain associated with tissue injury and infections, results from the heightened sensitivity of the peripheral terminals of nociceptor sensory neurons in response to exposure to inflammatory mediators. Targeting immune-derived inflammatory ligands, like prostaglandin E2, has been effective in alleviating inflammatory pain. However, the diversity of immune cells and the vast array of ligands they produce make it challenging to systematically map all neuroimmune pathways that contribute to inflammatory pain. Here, we constructed a comprehensive and updatable database of receptor-ligand pairs and complemented it with single-cell transcriptomics of immune cells and sensory neurons in three distinct inflammatory pain conditions, to generate injury-specific neuroimmune interactomes. We identified cell-type-specific neuroimmune axes that are common, as well as unique, to different injury types. This approach successfully predicts neuroimmune pathways with established roles in inflammatory pain as well as ones not previously described. We found that thrombospondin-1 produced by myeloid cells in all three conditions, is a negative regulator of nociceptor sensitization, revealing a non-canonical role of immune ligands as an endogenous reducer of peripheral sensitization. This computational platform lays the groundwork to identify novel mechanisms of immune-mediated peripheral sensitization and the specific disease contexts in which they act.

## Main Text

Inflammatory pain is caused by the sensitization of pain-triggering sensory neurons, nociceptors, by immune ligands, making the neuroimmune axis an attractive therapeutic target ^1, 2^. To delineate bidirectional neuroimmune interactions in diverse inflammatory pain conditions we performed single-cell transcriptomics of skin immune cells in three pain models: 1. **skin incision**, 2. **UV exposure**, and 3. **zymosan injection** and combined this with a single-nucleus sequencing (single-nuc seq) dataset of DRG neurons. We applied these data to a comprehensive receptor-ligand database which we compiled by combining structured interaction data with literature mining to predict neuroimmune interactions. This approach predicted a repertoire of known as well as novel neuroimmune interactions for each pain condition. Interestingly, each inflammatory pain model exhibited a significantly different immune cell state signature with a unique repertoire of potential receptor-ligand interactions between immune cells and sensory neurons. Immune-derived thrombospondin-1 (TSP-1) binding to neuronal CD47, was one of the novel neuroimmune axes found in all three conditions. TSP-1 attenuated prostaglandin E2 (PGE2)-induced peripheral sensitization in nociceptors, indicating that immune signaling can reduce, as well as generate, pain hypersensitivity. The data illustrates that different inflammatory conditions give rise to unique neuroimmune interactomes, and that unbiased mapping can aid discovery of novel pain mechanisms.

First, we assessed pain dynamics after skin incision, which models post-operative tissue injury, UV exposure, to model burn injury, and zymosan injection, to mimic infection-induced inflammatory pain. Mice subjected to either the skin incision or zymosan models developed significant hypersensitivity to noxious heat within 4 hours, which resolved by 48 and 24 hours, respectively. In contrast, heat hypersensitivity increased more slowly in the UV-exposed model and with no recovery at 48 hours (Fig 1A). This temporal dynamics of the onset and recovery suggests that these three pain conditions are distinct and may potentially be associated with varied immune responses. To determine if the differences in behavioral phenotype are caused by distinct neuroimmune interactions, we defined cell-type-specific neuroimmune interactions associated with these three pain conditions. We isolated immune cells at the time of maximally increased post-injury hypersensitivity, which was at 4 hours, 24 hours, and 48 hours for zymosan, skin incision and UV burn, respectively and performed single-cell sequencing of CD45+ immune cells from the inflamed and contralateral healthy skin (Fig 1B). To classify cell types in all conditions, we performed UMAP dimensionality reduction^3^ (Fig 1C). Cells from each biological replicate showed minimal batch-to-batch variation (Fig S1A). We then identified marker genes that define each cluster (Fig 1D and S1B). Dermal macrophages were identified by expression of *Csf1r, Cd163*, and *Mrc1* ^4^. Recruited macrophages (RM) showed an increased proportion in all injured states compared to healthy control skin. RMs also expressed reparative genes such as *F13a1, Selenop, Mrc1* and *Fn1*, suggesting an adaptive response to the injury. Three additional clusters displayed macrophage signatures (*Csf1r* and *Lyz2*) but could not be functionally defined by their transcriptional signatures, which we annotated as Macs 1, Macs 2, and Macs 3. Macs 1 was the inflammatory cluster, based on highest expression of *Il1b and Cxcl1*. Both Macs2 and Mac3 expressed tissue-resident markers, such as *Cd163* and *Mrc1*. The Macs2 cluster also expressed *Fn1*, like the RMs, suggesting that they while likely derived from infiltrating macrophages are in the process of acquiring a dermal macrophage profile (Fig 1C and 1D). We also identified two dendritic cell clusters, DCs 1 and DCs 2, based on expression of H2-Ab1. DCs 2 showed a higher expression of *Il1b*, highlighting its inflammatory characteristics. Finally, neutrophils (*S100a9* and *Csf3r*), Langerhans cells (*Csf1r* and *Cd74*), T cells (*Thy1* and *Icos*), Mast cells (*Mcpt4*) and Keratinocytes (*Krtdap*) were also clearly identified in the skin (Fig 1C and 1D). Single cell transcriptomics of skin captures, therefore, a broad spectrum of functionally diverse skin immune cells.

**Figure 1.**
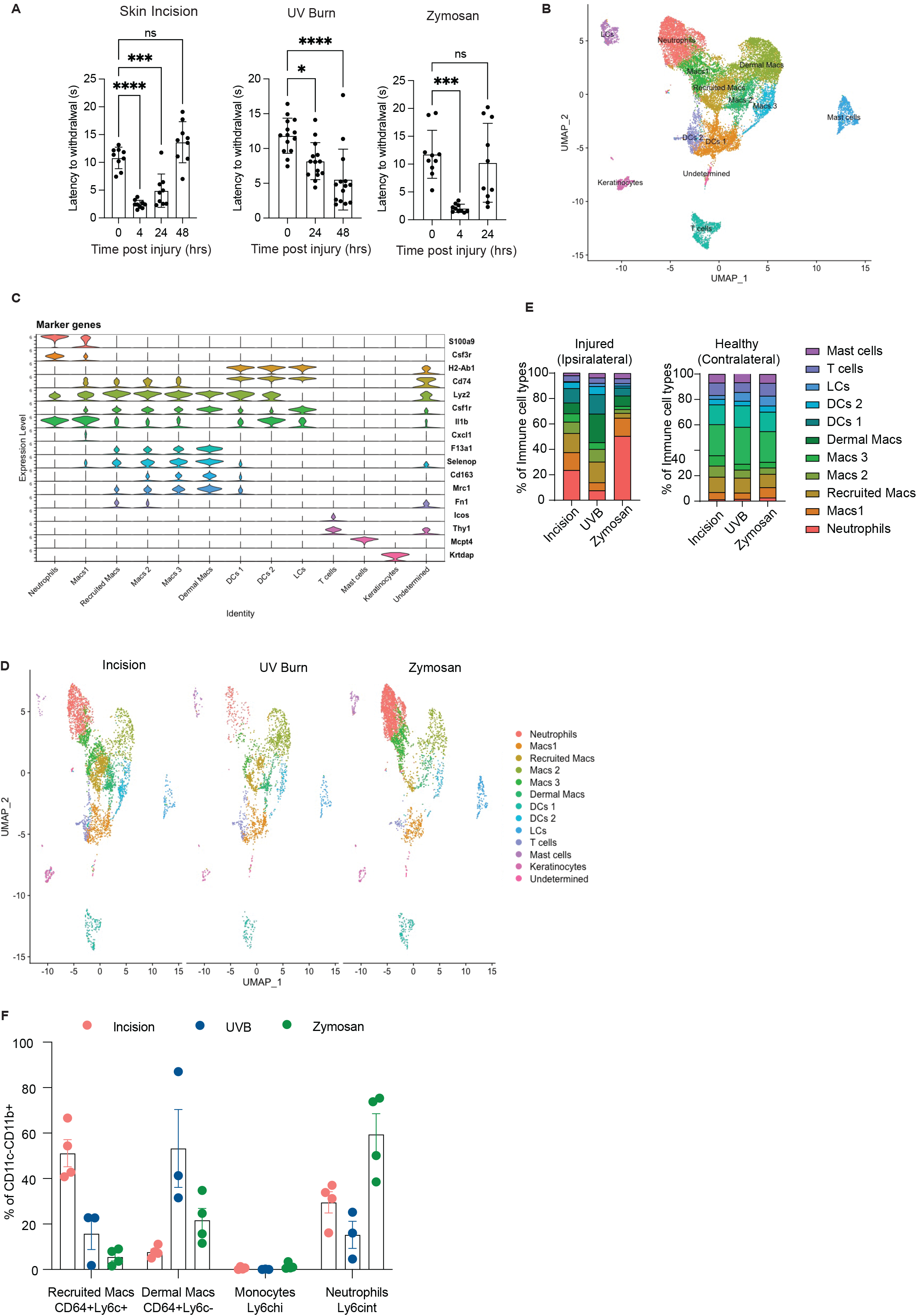
Distinct behavior and immune profiles in distinct inflammatory pain conditions. (A) Heat hypersensitivity in injured paws measured by the latency to react in the Hargreaves assay in WT mice at given times following skin incision, UV burn, and zymosan injection. p values calculated using oneway ANOVA, Tukey’s multiple comparison test. Both male and female mice were included. (B) A UMAP plot of merged single-cell RNA seq data from all the injured and contralateral healthy skin samples captures the heterogeneity of immune cells in the skin. (C) Violin plot showing expression of the specific marker genes used to assign an identity to the cell clusters. (D) UMAP plot of single-cell RNA seq data from immune cells in the paw skin after skin incision, UV burn, and zymosan injection (E) Proportions of immune cell populations in healthy contralateral paw skin were consistent between all models. (F) Immune cells isolated from injured or healthy paw skin stained for myeloid cell markers. Quantification of myeloid cell populations gated on live CD45+ CD11b+ CD11c-cells.

We compared the immune landscape of the three inflammatory pain conditions and detected an increase in neutrophils and RM populations in the skin incision and zymosan models compared to healthy controls (Fig 1D, 1E and Supp Fig 1B). In contrast, UV exposure only induced a moderate increase in these cell types. We also observed a significant contraction of the regulatory dermal macrophage population in both skin incision and zymosan-treated skin but not in the UV burn (Fig 1D, 1E and Supp Fig 1B). The immune populations in the contralateral healthy skin were comparable for all three injuries (Fig 1E and Supp Fig 1C). The cellular composition of the myeloid cells identified by the transcriptional profiling mirrored exactly that from flow cytometry analysis (Fig 1F and Supp Fig 2A). This data reveal that the three inflammatory pain conditions exhibit different immune cell signatures, and that single-cell transcriptomics provides detailed cell-type- and injury-specific gene expression profiles, which we could exploit to predict potential interactions with different sensory neuronal subsets.

To identify the transcriptional expression of receptors on DRG neurons complementary to ligands produced by the immune cells, we utilized a previously published dataset of single-nuc seq of mouse DRG neurons, from our lab ^5^. This dataset consists of nine subsets of sensory neurons, including peptidergic, non-peptidergic nociceptors, cLTMR, and Nefh+ neurons (Supp Fig 3A). We used this naïve mouse dataset to complement the transcriptomics of immune cells in inflammatory pain models, since an acute CFA-induced inflammation did not produce a significant transcriptional change in DRG neurons in this study ^5^. A differential expression analysis of all receptor genes (Supplementary data file 1) on all the DRG neurons revealed a unique receptor repertoire in the different subsets of sensory neurons (Fig 2A and Supp Fig 3B). Many of the differentially expressed genes were immune receptors, suggesting that nociceptors are poised to respond to multiple distinct immune signals.

**Figure 2.**
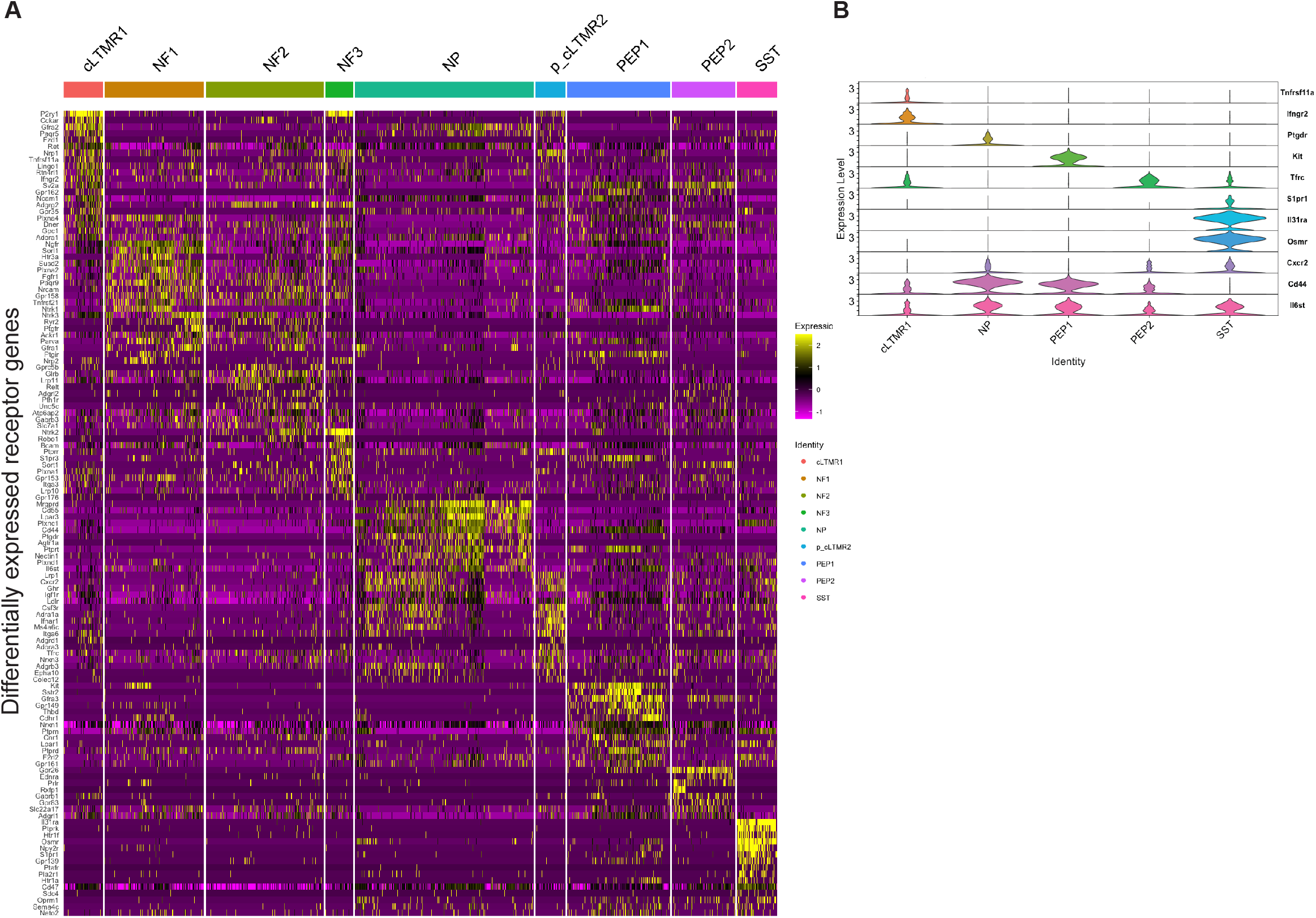
DRG neurons exhibit subtype-specific receptor expression profiles for immune ligands. (A) FindMarker analysis on receptor genes expressed by DRG neurons, from naïve female single-nuc RNA sequencing data, from Renthal et. al., (2020). Heatmaps show significantly (p<0.05) differentially expressed receptor genes neurons with log2FC >0.8 (B) Violin plot showing normalized expression of specific receptor genes in given DRG subtypes.

*Ifngr2*, which encodes the receptor for the type 1 cytokine IFNγ, was enriched in cLTMR neurons, suggesting that cLTMRs preferentially respond to IFNγ-producing cells such as CD8 T cells and NK cells (Fig 2B). On the other hand, Kit expression was only found on the peptidergic 1 population (Fig 2B). Kit binds to CD117, which is highly expressed by mast cells, a prominent skin-resident immune cell ^6^. Enriched Kit expression in peptidergic nociceptors implicates a potential role in type 2 immune responses. Peptidergic nociceptors amplify allergic immunity ^7, 8, 9^., and Kit-CD117 neuroimmune interactions could participate in this process. Furthermore, Osmr, the receptor for Oncostatin M (OSM) was specifically expressed by somatostatin (SST) neurons (Fig 2B), consistent with their role in OSM-mediated pruritus^10^. A small proportion of cLTMRs also uniquely expressed *Tnfrsf11a* that encodes RANKL (Fig 2B). Rankl is reported to play a role in peripheral sensitization ^11^ but cell type specificity was not identified in that study. The receptor expression analysis unveils that the neuronal cell type expressing RANKL could contribute to pain hypersensitivity. We also found that several receptors, traditionally studied for their function in immune cells, were also expressed by sensory neurons. For example, Cd44 and S1pr1 have an established function in immune cell extravasation and egress ^12^, and Cd44 was broadly expressed by C-fibers, while S1pr1 was highly expressed in SST neurons compared to other neurons (Fig 2B).

To more broadly define the relationship between immune cells and neurons in the three inflammatory models, we considered three possible mechanisms enabling crosstalk between these two cell types (Fig 3A). First, ligands secreted or expressed on the cell surface of immune cells may physically interact with cell surface receptors expressed on neurons. Second, metabolites whose production and secretion are controlled by enzymes in immune cells may interact with cell surface receptors in neurons, creating indirect interactions between immune cell enzymes and receptors. Finally, we also considered direct or indirect interactions of immune cell proteins with ion channels expressed on the cell surface of neurons. To assemble this interaction space, we used the Integrated Network and Dynamical Reasoning Assembler (INDRA) to draw on existing cell-to-cell interaction databases and combine these with interactions extracted by text mining of the scientific literature (Fig 3B)^13, 14^.

**Figure 3.**
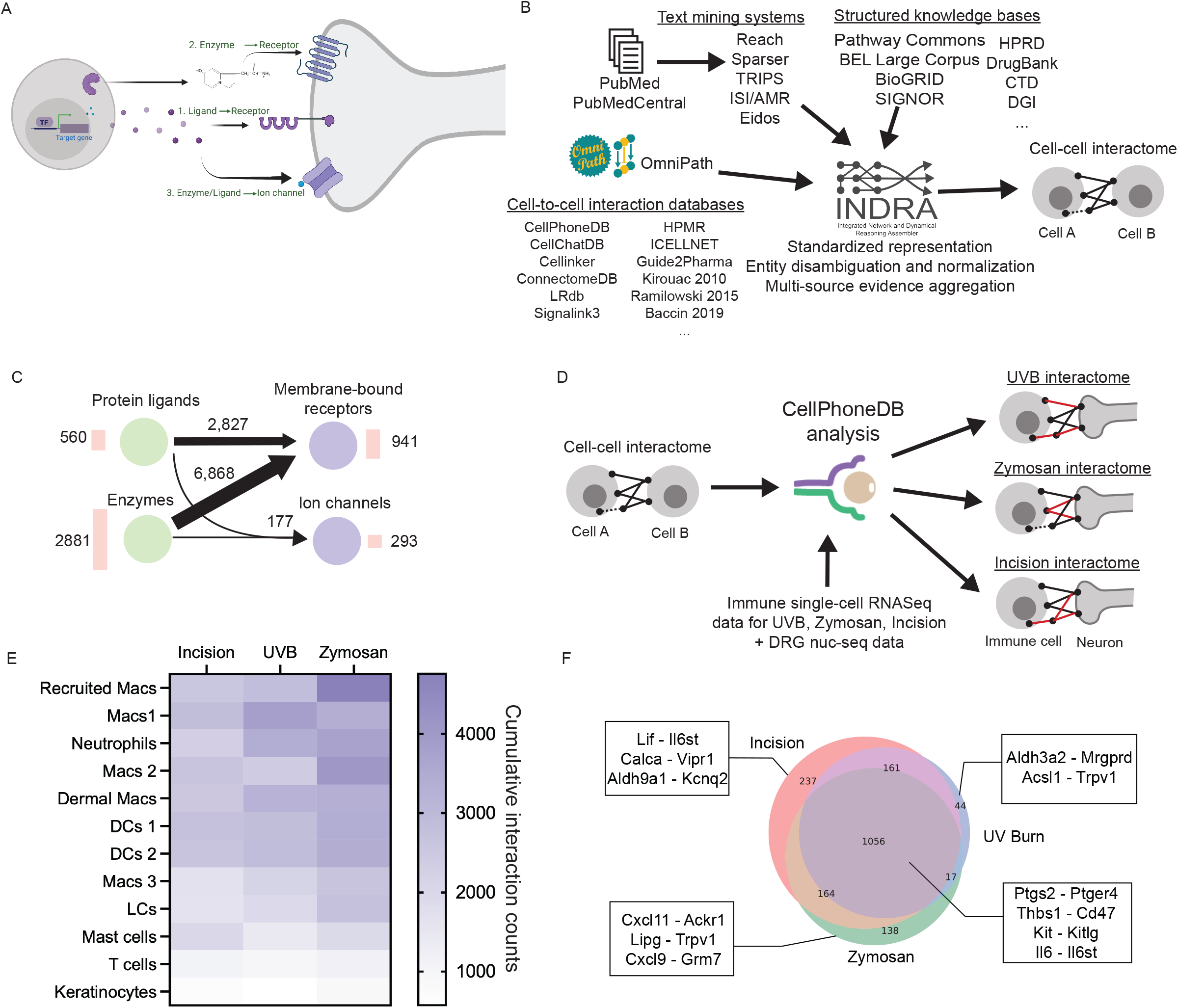
Construction of updatable and comprehensive neuroimmune interactome. (A) Schematic showing the types of intercellular communication considered between immune cells and neurons for constructing the interactome. (B) Overview of the computational pipeline for constructing the neuroimmune interactome. INDRA processes publications through multiple text mining systems and combines their output with structured knowledge bases integrated with INDRA directly, as well as the content of cell-to-cell interaction databases obtained via OmniPath. INDRA standardizes and aggregates evidence across all its sources and creates an assembled cell-cell interactome. (C) Summary of the interactome. Circles represent types of proteins: ligands, enzymes, membrane-bound receptors, and ion channels. The number of each protein type in the interactome is shown next to the circle. Arrows between circles show the number of distinct interactions among the corresponding protein type, with thicker arrows corresponding to a larger number of interactions. (D) Overview of the condition-specific interactome construction. The general cell-cell interactome was provided as input to CellPhoneDB together with data from immune cells for each of the three pain conditions: UVB, Zymosan or incision and combined with naïve DRG neuron data. This results in three condition-specific neuroimmune interactomes. (E) Heatmap of the total number of significant bidirectional interactions between immune cells and neurons calculated for each injury. The number of neuroimmune interactions was calculated, omitting immune-immune and neuronneurons interactions. Color saturation represents a higher number of neuroimmune interactions. (F) Venn diagram depicting the number of shared and unique significant neuroimmune interactions for each inflammatory pain condition.

First, we defined that set of ligands, receptors, enzymes, and ion channels to consider in the interactome. To construct the set of ligands and receptors, we used the CellPhoneDB interactome as a reference^15^. For each interaction, we classified one of the interactors as the receptor and the other as the ligand, based on a set of consensus direction information standardized from 10 sources by OmniPath^16, 17^ (see Methods). We constructed the enzyme list based on the ExPASy Enzyme database^18^, capturing human proteins known to have enzymatic activity, excluding kinases and phosphatases because they are intermediaries in signaling pathways and not directly involved in intercellular interactions. Finally, we used a list of ion channels curated in the NIH Illuminating the Druggable Genome program (https://druggablegenome.net/). Together these analyses yielded 941 receptors (Supplementary data file 1), 560 ligands (Supplementary data file 2), 2,881 enzymes (Supplementary data file 3), and 293 ion channels (Supplementary data file 4), (Fig 3C).

From the ligand, receptor, enzyme, and ion channel protein sets, we assembled all potential interactions (Supplementary Data File 1). Ligand-receptor interactions were obtained from OmniPath, which aggregates more than a dozen cell-to-cell interaction databases. We aligned these interactions with the text mining and structured knowledge base evidence collected by INDRA, to obtain 2,827 distinct interactions (Figure 3C). We used the Pathway Commons database^19^ to determine each enzyme as well as their products, and then used INDRA to find interactions between these products and receptors. Where an enzyme’s product interacted with a receptor, we created an interaction (6,868 in total, Figure 3C), while maintaining the underlying relations between enzymes, products, and receptors for traceability. Finally, we used INDRA to find direct or indirect effects on ion channels from protein ligands or enzyme products and added an additional 177 such interactions (Figure 3C). The interactome thus contains 9,872 unique interactions in total.

This initial interactome defines all possible interactions between immune cells and neurons and is not customized to specific cell types or contextualized based on specific data. It serves though, as a versatile toolkit for other analyses. Given INDRA’s ability to standardize and align representations of interactions from structured sources and text mining, the interactome is tied directly to the scientific literature ^13^. In addition, since INDRA can process new literature as it appears, the interactome can be updated. We created a web-browsable version of the interactome where each interaction appears as a heading that can be opened to examine evidence from publications supporting a given interaction (Supp Fig 4A). We used this interactome to then identify potential significant neuroimmune interactions for the three specific inflammatory pain conditions (Supplementary Data File 1).

CellPhoneDB was used to identify statistically significant interactions between each immune and neuronal cell type based on receptor-ligand expression profiles for each of the three pain conditions (Fig 3D) ^15^. We merged the three contralateral datasets to identify receptor-ligand expression in healthy conditions. The single-cell transcription data from each condition was combined individually with the healthy DRG neuron dataset ^5^. All sensory neuronal subtypes were considered in the cell-cell interaction analysis. The analysis produced a network of all bidirectional interactions that can occur between the distinct neuronal and immune cell types in healthy and inflammatory pain conditions (Supplementary Data File 2-5). The interactome network also includes immune-immune as well as neuron-neuron interactions (Supp Fig 4B). To specifically identify pathways potentially relevant to inflammatory pain, we only focused on immune-neuron interactions and the significant cumulative interactions of all neurons (cLTMR, NP, Pep1, Pep2, SST, NF1, NF2, NF3 and p_cLTRM2) with each immune cell type were calculated (Fig 3E). RM and the inflammatory Macs 1 cluster had the highest cumulative interactions with sensory neurons, when compared to the healthy skin interactome which was generated by merging data from all contralateral control samples (Fig 3E).

The neuroimmune interactions predicted for each of the three models were then compared. Several of these with an established role in inflammatory pain, such as, Ptgs2-Ptger4, Kit-Kitlg, and Il6-Il6st, were shared between all models representing common inflammatory pain-promoting pathways (Fig 3F and Fig 4). The analysis also predicted interactions unique to individual inflammatory pain models. Lif – Il6st, Cxcl11 – Ackr1, and Aldh3a2 – Mrgprd, which were unique to incision, zymosan, and UVB, respectively (Fig 3F). In addition to these known pathways, the analysis also predicted novel neuroimmune axes in inflammatory pain, such as Spp1-S1pr1, Spp1-Cd44, Thbs1-CD47, Thbs1-Lrp1, and Ccl18-Ackr1 (Fig 3F). This suggests that cell-type-specific transcriptional expression data can be used as a platform for predicting physiologically relevant pain-sensitizing pathways from immune cells.

**Figure 4.**
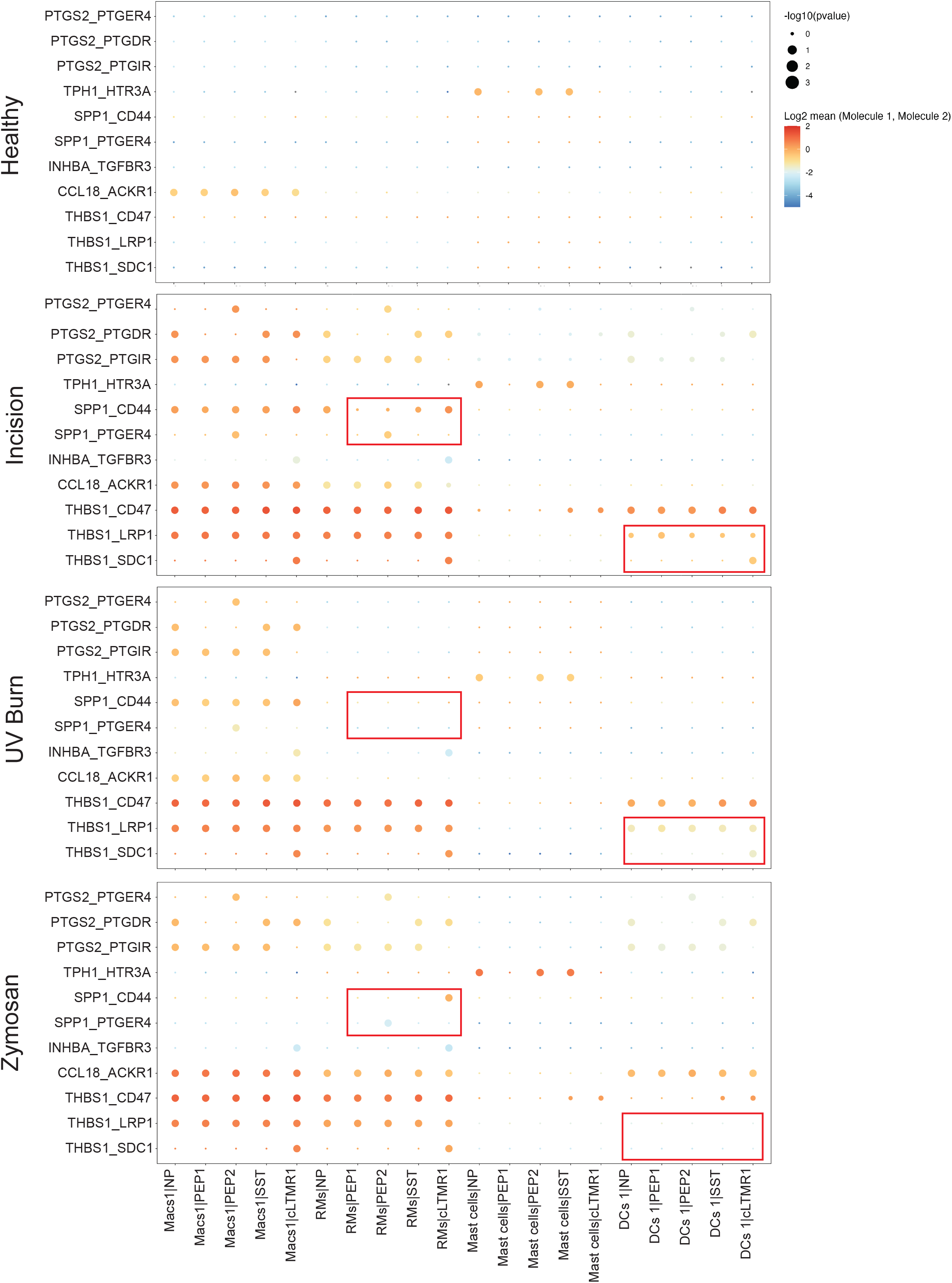
Neuroimmune receptor-ligand interactions occur in an injury- and cell-type-specific manner. Dot plots showing interaction strengths between neuron-immune cell pairs for defined receptor-ligand pairs. The strength is determined by the statistical method in Cellphonedb. The y-axis shows the cell-cell pairs and the x-axis ligand-receptor pairs. The size of the dot shows the significance of the interaction, which is determined by how specific this interaction is to the specific cell-cell pair. The scale of the plots is determined by the expression levels of receptor and ligand genes in each cell type. The red boxes outline cell-type specific interactions that differ between injury types.

The single-cell transcriptomics also allowed us to identify cell-type-specific ligand-receptor interactions. For instance, Ptgs2 expressed by inflammatory Macs (Macs1) and RMs were likely to interact with the EP4 receptor on Pep2 neurons, while on NP, SST, and cLTMR neurons, the prostaglandin D2 receptor (Ptgdr) was more likely to be engaged (Fig 4). Osteopontin (OPN), encoded by *SPP1*, was predicted to enable an interaction between RMs and neurons only in the skin incision model, while the Macs1 population could interact with neurons via OPN in the UV burn model. The Thbs1-Lrp1 axis was engaged by DCs only in skin incision and UV burn, but not the Zymosan model (Fig 4).

One interaction pathway shared by all three pain conditions is Thbs1-CD47 (Fig 3F). *Thbs1* encodes thrombospondin-1 (TSP-1) which binds to several known receptors, including a2d1, LRP1, CD47, and a2b3 integrin ^20^. TSP-1 is traditionally studied in the context of platelet activation, where it acts on CD47 on platelets to trigger Gi-coupled GPCR signaling attenuating PKA activation ^21, 22^. We detected an upregulation of Thbs1 in immune cells during inflammatory pain hypersensitivity (Fig 5A) and found that the CD47 receptor, together with other TSP1 receptors, is highly expressed by mouse DRG neurons (Fig 5B and 5C). In nociceptors, PKA activation triggers several sensitization pathways ^23, 24, 25^ by directly phosphorylating TRPV1 and Nav1.8 channels leading to a reduced activation threshold ^26^. We hypothesized that TSP-1 may exert an effect on DRG neurons through CD47, inhibiting the PKA-mediated Trpv1 sensitization. We tested this using a capsaicin-induced peripheral sensitization assay in cultured mouse DRG neurons. Capsaicin-responsive (Trpv1+) nociceptor neurons were exposed to PGE2, and intracellular calcium levels measured with the calcium indicator, Fura-2. The amplitude of the Ca^2+^ influx triggered by low-dose capsaicin was significantly higher in DRG neurons treated with PGE2 (5D and 5E), consistent with a PGE2-induced Trpv1 sensitization ^27^. The majority of DRG neurons did not respond to low-dose (100nM) capsaicin, without prior sensitization (Fig 5D, *black trace*). We only observed the PGE2-induced Trpv1 sensitization in DRG neurons from male mice, female DRG neurons showed no significant increase upon PGE2 exposure (Sup fig 5A). When DRG neurons were co-treated with TSP-1 we observed a significant attenuation of the PGE2-induced sensitization (5D and 5E). This validated the predicted interaction of TSP-1 with DRG neurons and highlights a non-canonical function of immune ligands, one that counteracts neuronal sensitization, and as an endogenous suppressor/modulator of inflammatory pain hypersensitivity.

**Figure 5.**
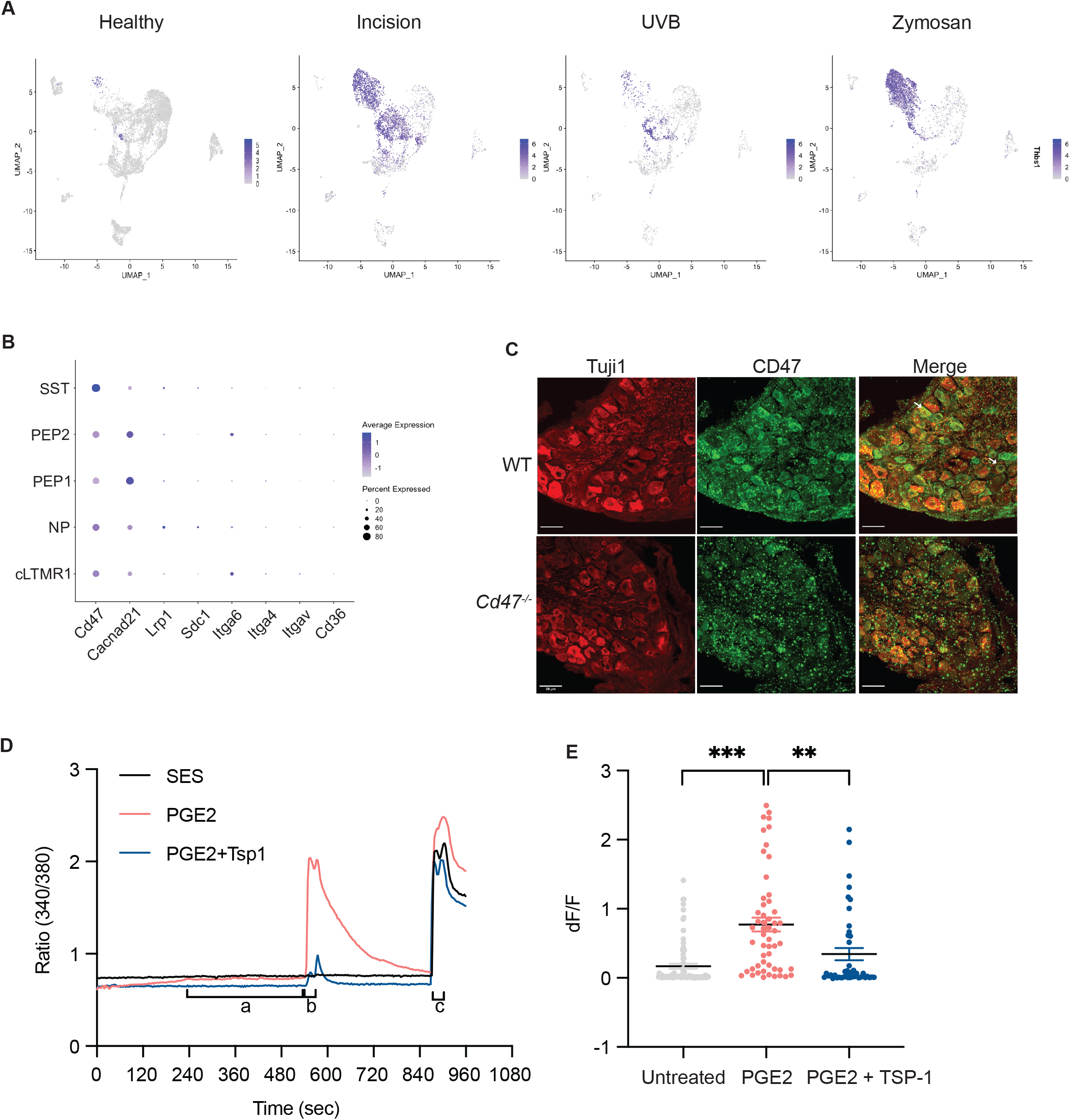
Thrombospondin-1 act on DRG neurons to inhibit PGE2-mediated sensitization. (A) U-MAP plot of normalized Thbs1 expression in healthy and injured skin immune cells. (B) Dot plot showing expression of known TSP-1 receptors in C-fiber DRG neuron subtypes. (C) CD47 (Green) expression on frozen WT or *Cd47*^-/-^ DRG neuron sections stained with Tuji1 (Red). (D) Fura-2-based calcium imaging was performed on cultured DRG neurons from healthy male mice. SES was used as the recording solution. Treatments were applied during live imaging using a gravity-based perfusion system. Cells were treated with (*1*) SES (black), 1uM PGE2 (Red) or 1uM PGE2 + 200ng/ml TSP-1 (Blue) for 7 min, (*2*) immediately followed by (a) 100nM Capsaicin (black), 100nM Capsaicin + 1uM PGE2 (Red) or 100nM Capsaicin + 1uM PGE2 + 200ng/ml TSP-1 (Blue) for 30 seconds. Cells were then washed with SES for 5 min followed by (*3*) treatment with 1uM Capsaicin for 30 seconds to reveal TRPV1+ neurons. Frames were captured every 3 seconds. Intensity traces of ratio of 340/380 are plotted. (E) dF/F was calculated as (F_1_-F_0_)/F_0_, where F_1_ is the peak response within 40 seconds of treatment *2*, F_0_ is the average of 10 seconds prior to treatment *2*. All TRPV1+ neurons are shown from one experiment consisting of three separate recordings for each condition. Unpaired t-test was performed, p < 0.05 was considered significant. Data is representative of three independent experiments.

These findings highlight the highly contextual nature of neuroimmune interactions^28^. While the three distinct models used in this study all showed inflammatory signatures compared to their healthy counterparts, the actual cellular and transcriptional makeup of the individual models is different. The distinct immune responses to different forms of tissue injury align with our understanding of different immune responses but calls for a closer look into how sensory neurons sense immune-derived cues, especially since neuronal subsets differentially express receptors for immune ligands. Several interactions were shared by all three inflammatory pain conditions and many of these are known to promote peripheral sensitization, however, we unexpectedly identified TSP-1, an immune ligand, as a novel suppressor of peripheral sensitization. TSP-1 is highly upregulated after tissue injury by myeloid cells and has a tissue reparative role. TSP-1 acts on endothelial cells to suppress vascularization and nitric oxide production, which promotes tissue repair ^22, 29^. In the central nervous system, TSP-1 is produced by astrocytes and promotes synaptogenesis ^30^. Our finding that TSP-1 inhibits PGE2-induced peripheral sensitization indicates a novel suppressive function for this ligand in the peripheral nervous system. While the endogenous opioid system is well studied for its role in inhibiting pain via actions on peripheral and central neurons that express the mu opioid receptor (MOR) ^31^, TSP-1 is a novel immune-derived inflammatory pain suppressor, directly acting on the peripheral terminals of sensory neurons in the periphery. Infiltrating macrophages that produce TSP-1 also secrete several inflammatory mediators such as PGE2, IL1β, and TNFα. TSP-1 is likely then to act in concert with that inflammatory signaling to finely modulate the overall pain sensitivity outcome. It is now evident that immune ligands present in the complex inflammatory milieu in skin inflammation exert opposing effects on primary sensory neurons, and that inflammatory pain is likely then, to be more complex than previously considered, with both drivers and suppressors.

## Acknowledgments

We would like to thank the Research Computing Core at BCH and the HMS Single Cell Core at HMS for their services and advice on experimental strategy and analysis. This study is funded by DARPA HR0011-19-2-0022 and NIH R35NS105076 to C.J.W. and Jane Coffin Childs Fund to A.J.

## Methods

### Mice

8-12-week-old C57BL/6J mice were obtained from the Jackson Laboratory (JAX:000664). Cd47 deficient mice were obtained from Jackson Laboratory (Jax:003173). Both Male and Female mice were used in all behavior experiments. Single-cell RNA sequencing was performed on female mice. All animal experiments were conducted according to institutional animal care and safety guidelines at Boston Children’s Hospital and Harvard Medical School.

### Inflammatory pain models

For each model, mice were randomly selected from the cage to receive the inflammatory stimulus. All mice in a cage received the same inflammatory stimulus and the person inducing the inflammation was different to the person testing for hypersensitivity.

#### Skin Incision

Mice were anesthetized by administration of 2.5% isoflurane. 3-5mm skin incision was made using a surgical sterile scalpel without cutting through the underneath muscle. Skin was sutured using 6-0 silk surgical suture (Ethicon, K889H) in an aseptic manner. Contralateral control was left untouched.

#### UV burn

Mice were anesthetized by administration of 2.5% isoflurane. Mouse hind paw was exposed to UV irradiation at an intensity of 1 J/cm2 for 1 min using a wavelength 305nm – 315nm fluorescent UV-B light source. Contralateral control was not exposed to UV.

#### Zymosan

Mice were anesthetized by administration of 2.5% isoflurane. 20ul of 5mg/ml zymosan (in saline) was injected into the plantar surface of the hind paw. Saline was injected into the contralateral control.

### Hargreaves

Mice were habituated to the Hargreave’s apparatus (IITC #390G) consisting of a glass floor heated to 30C and a plexiglass chamber for 2 days prior to testing, 1 hour per day. On the day of assessment mice were habituated for another 1 hour before the test. A focused radiant heat light (15% intensity) source was focused on the plantar surface of the left paw of mice and a ramping heat stimulus was applied until a paw withdrawal was recorded. Readings were averaged from two trials. Blinded testing of mice with paw incision and zymosan injection was not possible since it was clear to see which paw was inflamed, but the operator was not told which foot the UV burn had been applied to, thus aiding blinding for those mice.

### Tissue processing of mouse plantar skin

The planter skin of the mouse hind paw was dissected, separating the muscle and collected into 1% BSA containing RPMI. The skin was minced using scissors into 1-2mm pieces. Liberase TM (Roche) was added to the media at a final concentration of 0.5 mg/ml. Tissue was digested at 37c while vortexing at 1000rpm for 90 min. The digested tissue was strained using a 100uM strainer to obtain a single-cell suspension used for flow cytometry and single cell transcriptomics.

### Flow Cytometry

Cells were washed with facs buffer and incubated with Fc block for 10 min on ice. Cells were then stained with mouse antibodies against mouse CD45-FITC (1:400), CD11b-ef780 (1:400), CD64-PE-594 (1:500), Ly6c-BV711 (1:1000), CD11c-APC (1:200) all from Biolegend, for 30 min on ice. Cells were washed and then resuspended with 3uM DAPI for used for analysis. All flow cytometry analysis was performed on BD Fortessa.

### Single-cell RNA sequencing of immune cells (inDrops)

For isolation of immune cells for single cell sequencing, cells were washed with facs buffer and incubated with Fc block for 10 min on ice. Cells were then stained with anti-mouse CD45-FITC antibody for 30 min at 4c. CD45+ DAPI neg cells were sorted to a 90% purity using BD ARIAII at Boston Children’s Hospital Flow cytometry core. Cells were collected into 0.5% BSA containing PBS without EDTA. Single cells suspensions were encapsulated into droplets at the singe cell sequencing core, Harvard Medical School. The RNA in each droplet was reverse transcribed using a unique oligonucleotide barcode for each nucleus as described previously^32^. After encapsulation and barcoding, 4 samples were pooled with 3,000 droplets per sample for library preparation (). Libraries were sequenced on an Illumina Nextseq 500 to a depth of 400 million single-end reads per ~12,000 droplets collected. Sequencing data was processed and mapped to the mouse genome GRCm38 using the pipeline described in https://github.com/indrops/indrops^32^. Count tables from each library were then processed as described below.

### Initial quality control, clustering and visualization of snRNA-seq

To be included for analysis, cells were required to contain counts of greater than 300 and fewer than 2800 unique genes, more than fewer than 5% of the counts derived from mitochondrial genes. The cells counts that met these criteria is as follows: Incision - 4286, Incision_contra - 3461, UVB - 2125, UVB_contra - 2110, Zymosan - 3787, Zymosan_contra - 2629. We used the Seurat package (4.1.0) in R to perform the clustering of these cells as previously described^33^. Counts were centered and scaled for each gene. SCT transform was used for normalization and regressing variable features. All the 12 sample datasets were then integrated before clustering. Cell clustering was performed using FindClusters() based on the top 40 principal components, with resolution at 0.6 for the initial clustering of all cells. For dimension reduction and visualization, Uniform Manifold Approximation and Projection (UMAP) coordinates were calculated in the PCA space by using the implemented function runUMAP() in Seurat.

### Analysis of DRG neuron Single nuc seq from Renthal et. al. (2020)

DRG neurons single nuc seq data was obtained from Renthal et. al.^5^ Neuronal cells from Naïve mice (male and female) were obtained by sub setting the Seurat object based on annotation. The counts were then scaled and normalized before combining them with the immune single cell seq data.

### Assembling the receptor-ligand database

INDRA 1.21.0 was used to assemble the interactome. The list of receptors was obtained from CellPhoneDB 2.1.7, using the *protein_generated.csv* file generated by CellPhoneDB’s *generate_proteins* function. OmniPath interactions were obtained on March 22, 2022, through INDRA’s OmniPath API, which uses the http://omnipathdb.org/interactions endpoint of the OmniPath web service to obtain the “ligrecextra” subset of interactions corresponding to ligand-receptor interactions. These interactions were then filtered for curation effort > 0, to ones containing human proteins only, and to the following OmniPath source identifiers corresponding to sources of cell-cell interaction information: “CellPhoneDB”, “Guide2Pharma”, “HPMR”, “ICELLNET”, “Kirouac2010”, “CellTalkDB”, “CellChatDB”, “connectomeDB2020”, “Ramilowski2015”, and “talklr”. The ligand list was derived by taking the “source” participant of OmniPath interactions with a well-defined consensus direction and excluding known receptors. The ion channel list was derived from a curated list of proteins from the Illuminating the Druggable Genome project at https://druggablegenome.net/, removing overlaps with any receptors and ligands. The enzyme list was obtained from Expasy at ftp://ftp.expasy.org/databases/enzyme/enzyme.dat via PyOBO (https://github.com/pyobo/pyobo) to extract proteins that belong to any enzyme class, then filtered out any proteins known to be kinases or phoshphatases based on lists maintained by INDRA. Ligand-receptor interactions were taken from the overall OmniPath interaction list by filtering to interactions containing one ligand and one receptor. Evidence from publications and further structured databases aligned with these interactions was then obtained via INDRA. Ligand-ion channel interactions were obtained from INDRA directly by filtering Complex and Activation INDRA Statement types and Statements containing one ligand and one ion channel per the gene lists above. Statements were also filtered to ones supported by structured databases or at least two supporting sentences from text mining. Interactions involving enzymes were obtained in two parts. First, Pathway Commons v12 data was obtained in SIF format from https://www.pathwaycommons.org/archives/PC2/v12/PathwayCommons12.Detailed.hgnc.sif.gz and filtered to controls-production-of interaction whose controller is in the list of enzymes. The set of products for each enzyme was determined from the collection of rows remaining after these filters. For each enzyme product, INDRA was then used to find Activation and Complex Statements in which the enzyme product interacts with a receptor or an ion channel. Finally, enzyme-receptor and enzyme-ion channel interactions were generated by connecting an enzyme to a receptor or ion channel if its product interacts with the given receptor or ion channel. Finally, the interactome was exported into a tabular format compatible with CellPhoneDB using UniProt IDs to identify interacting proteins. Source code to generate the interactome and the interactome gene lists used in this study is available at: https://github.com/indralab/neuroimmune_interactome.

### Identification of significant receptor-ligand interactions for cell-cell pairs

Normalized counts from Indrop seq of immune cells from each injury model was merged individually with the neuron counts object. One merged Seurat object was obtained from each condition: Incision, UVB and Zymosan. Contralateral controls for all three models were combined and termed as Healthy. CellphoneDB statistical methods was applied to the resulting four neuroimmune single cell RNA counts files. The user-defined receptor-ligand database was used (Supplementary Data File 1). The significant interactions in Healthy (Supplementary data file 1), Incision (Supplementary Data File 2), UV Burn (Supplementary data file 3) and Zymosan (Supplementary data file 4) were obtained based on the statistical method in CellphoneDB.

### Isolation and in vitro culture of DRG neurons

Lumbar and Thoracic DRGs were obtained from mice and collected in DMEM containing FCS and Penicillin and Streptomycin (DMEM). DRGs were digested in Collagenase A (Roche, 5mg/ml) and Dispase II (Roche, 1mg/ml) for 70 min. Digested DRGs were triturated using large, medium and small sized polished glass pipettes in DMEM containing DNase. The cells were resuspended in DMEM and overlayed on 10% BSA solution. The bilayer was centrifuged for 12 min at 1000g at reduced acceleration and deceleration. The top two layers were discarded and the cell pellet was collected. For Ca imaging, cells were cultured at the center of a PDL (500ug/ml) and Laminin (5mg/ml) coated 35mm dish in Neurobasal A Media (Life Technologies, 10888-022) containing 2% B-27, Penicillin, Streptomycin, 10 μM arabinocytidine (Sigma) and GDNF (Sigma Aldrich, SRP3200). 5000-8000 cells were plated in 50-80ul drop and replenished with 2ml of media per dish and incubated at 37c and 5% CO2 for 48hrs before Calcium imaging experiments.

### Calcium imaging of DRG neurons

After 48 hours in culture, DRG neurons were loaded with 4ug/ml of Fura2-AM (Invitrogen) by incubating for 50 min at RT. Cell were then washed with SES twice and left in 2ml of SES to be used as recording solution. Live imaging was performed on Nikon Eclipse Ti microscope (Nikon, Melville, NY) with standard 340- and 380-nm filters controlled by a Ludl Mac6000 shutter using Nikon Elements software. Frames were recorded every 3 seconds. All imaging was performed at room temperature. Cell treatments were performed using a gravity-based perfusion system. Cells were simply treated with SES for 2 min to get stable images and remove perfusion related artifacts. Then, cells were exposed to with freshly prepared PGE2 (1μM) +/-TSP-1 (200ng/ml) for 7 min, followed by the application of a low concentration of capsaicin (0.1 μM) for 30 sec with or without PGE2 or TSP-1. After a 5 min perfusion of SES, a high concentration of capsaicin (1 μM) was applied at 24 for 30 s, to identify all capsaicin-sensitive Trpv1+ neurons.

### Immunohistochemistry

DRGs were collected in PBS and then fixed in 4% PFA on a shaker at 4C for 30min. After fixation, DRGs were moved to 30% sucrose overnight then mounted in OCT and frozen. DRGs were sectioned at 20um, mounted onto superfrost plus microscope slides and stored at −20C. Slides were thawed for 30 minutes then washed with PBS for 15 minutes. Slides were incubated in blocking buffer for 2h at RT (10% Donkey Serum, 0.4% Triton-X, 0.05% Tween 20, 1% BSA) and then incubated in primary antibody overnight at 4C (Rb anti-Tubb3 Sigma Aldrich T2200 1:500, Goat anti-CD47 RnD Systems AF1866 1:200). Slides are then washed 3x in PBS and incubated in secondary antibody at RT for 2h (Donkey anti-Rb Cy3 Jackson ImmunoResearch 711-165-152 1:500, Donkey anti-Goat 647 Thermo Fischer A-21447 1:500). Slides were finally washed 3x with PBS and mounted with Prolong anti-fade DAPI medium. Stained slides were imaged on a Leica SP8 confocal microscope with a 63x oil objective. Z-stacks spanning the tissue were taken and 4 adjacent fields were tiled. ImageJ was used to obtain the maximum intensity projection.

### Data and Statistical Analysis

Statistical analysis, including animal numbers (n) and p values, are included in the figure legends. Statistical analysis was performed using Graphpad Prism 9. For CellphoneDB package the statistical method was used, which considers the p.value for interactions that is defined by the enrichment of the interacting ligand-receptor pair in each of the interacting pairs of cell types. The p-value for significant interactions was <0.05. CellphoneDB was used on Python 2.7. All single-cell sequencing analysis was performed on RStudio version 1.3.959.

---

**Supplementary Figure 1.**
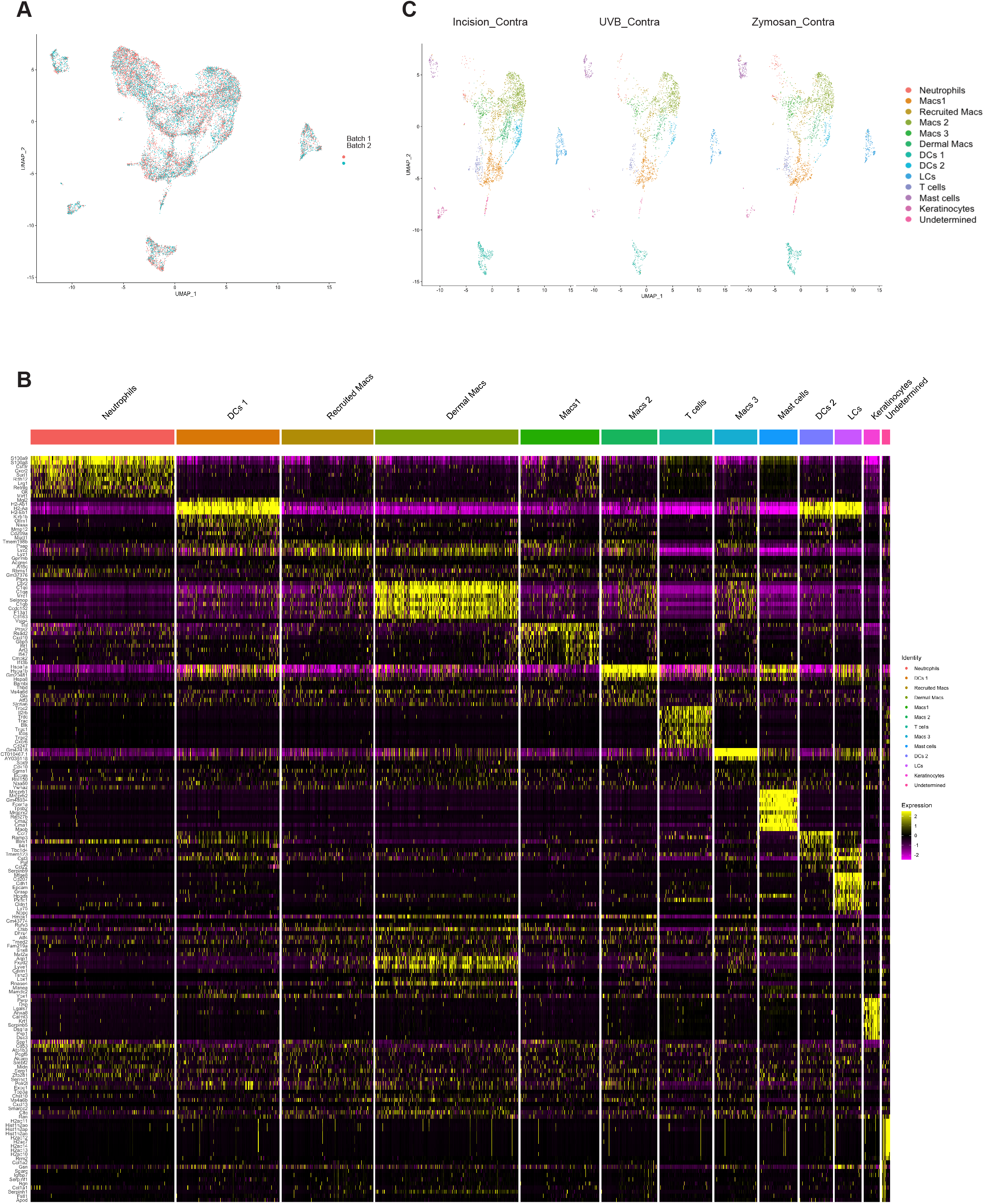
Single-cell RNA sequencing analysis of immune cells. (A) UMAP plot showing cells from single-cell sequencing from the two batches of sequencing experiments. (B) Heatmap showing top 40 marker genes for each cell cluster (C) UMAP plot of single-cell RNA seq data from immune cells in the contralateral healthy paw skin after skin incision, UV burn, and zymosan injection.

**Supplementary Figure 2.**
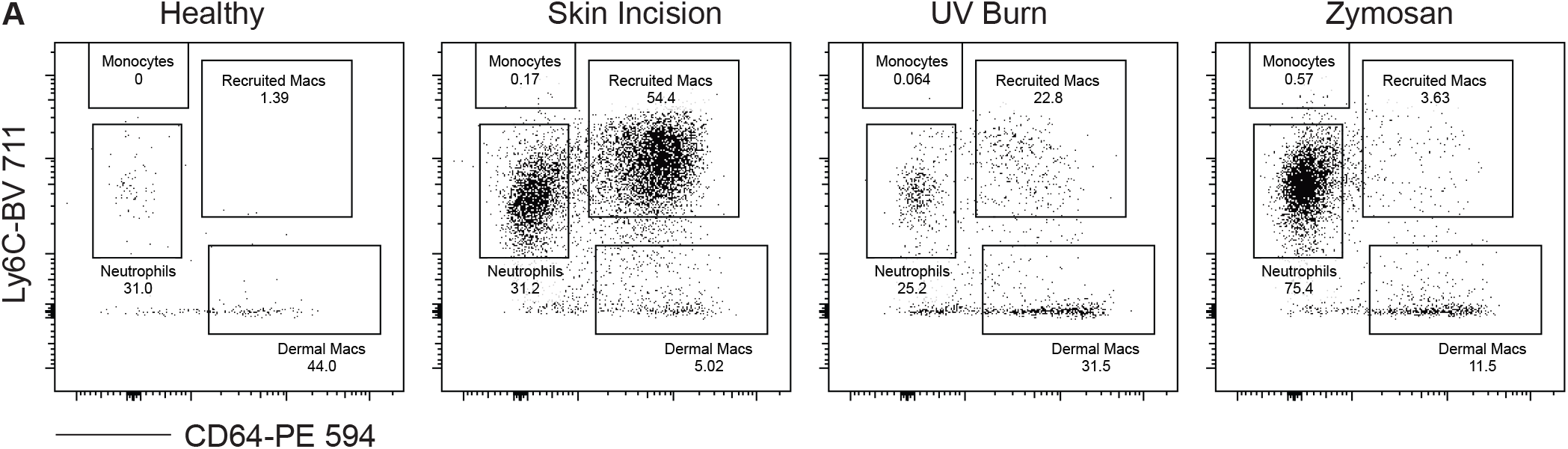
Myeloid cell populations in different inflammatory pain injuries. Immune cells were isolated from injured or healthy paw skin and stained for various myeloid cell markers. (A) Representative flow plots are gated on Live CD45+ CD11b+ CD11c-cells.

**Supplementary Figure 3.**
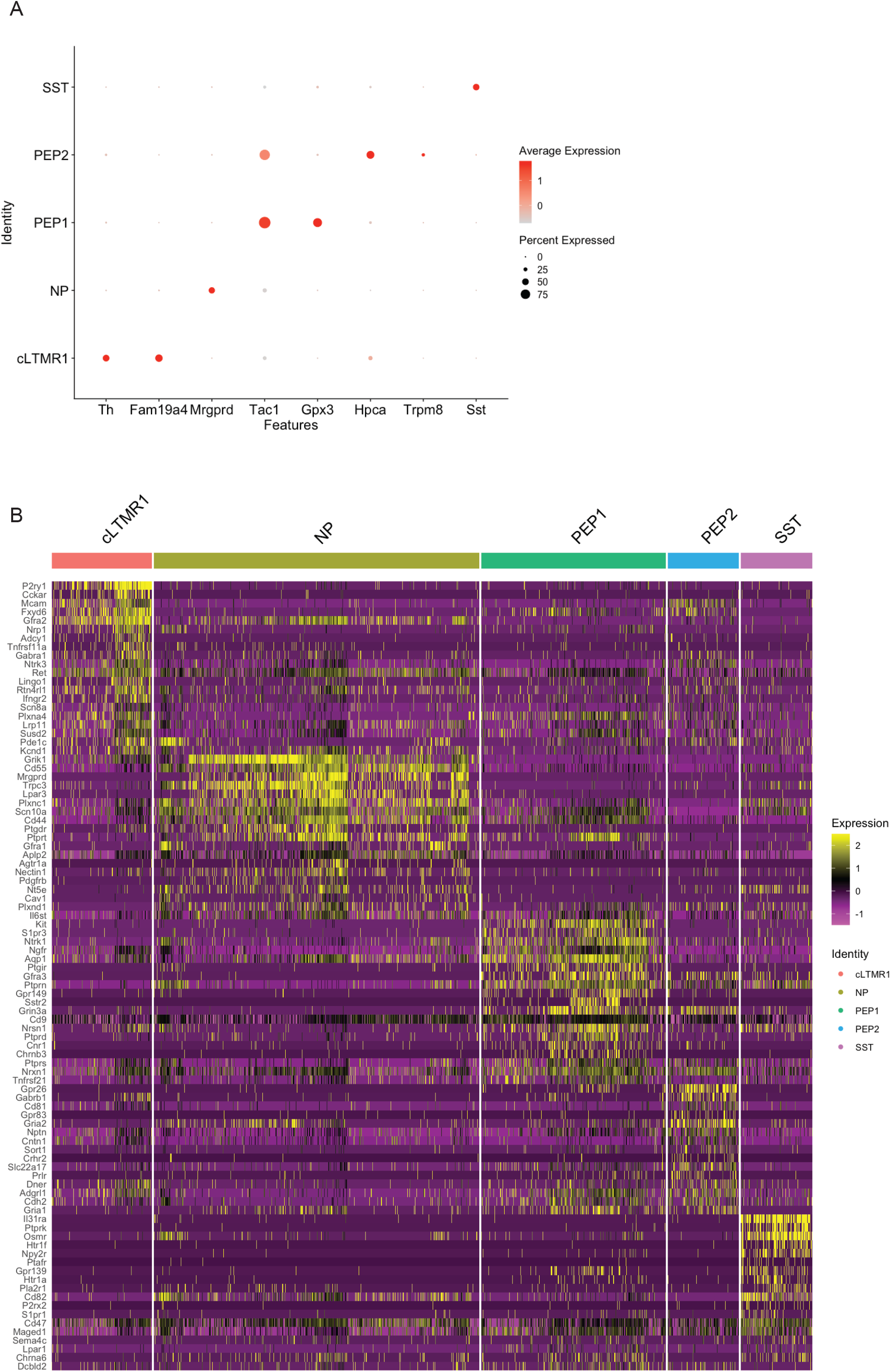
Receptor expression profile in male DRG neurons. (A) Dot plot showing marker genes used to annotate given DRG subtypes. (B) Differential expression of receptor genes in PEP1, NP, SST, and cLTMR cells from male DRG neurons.

**Supplementary Figure 4.**
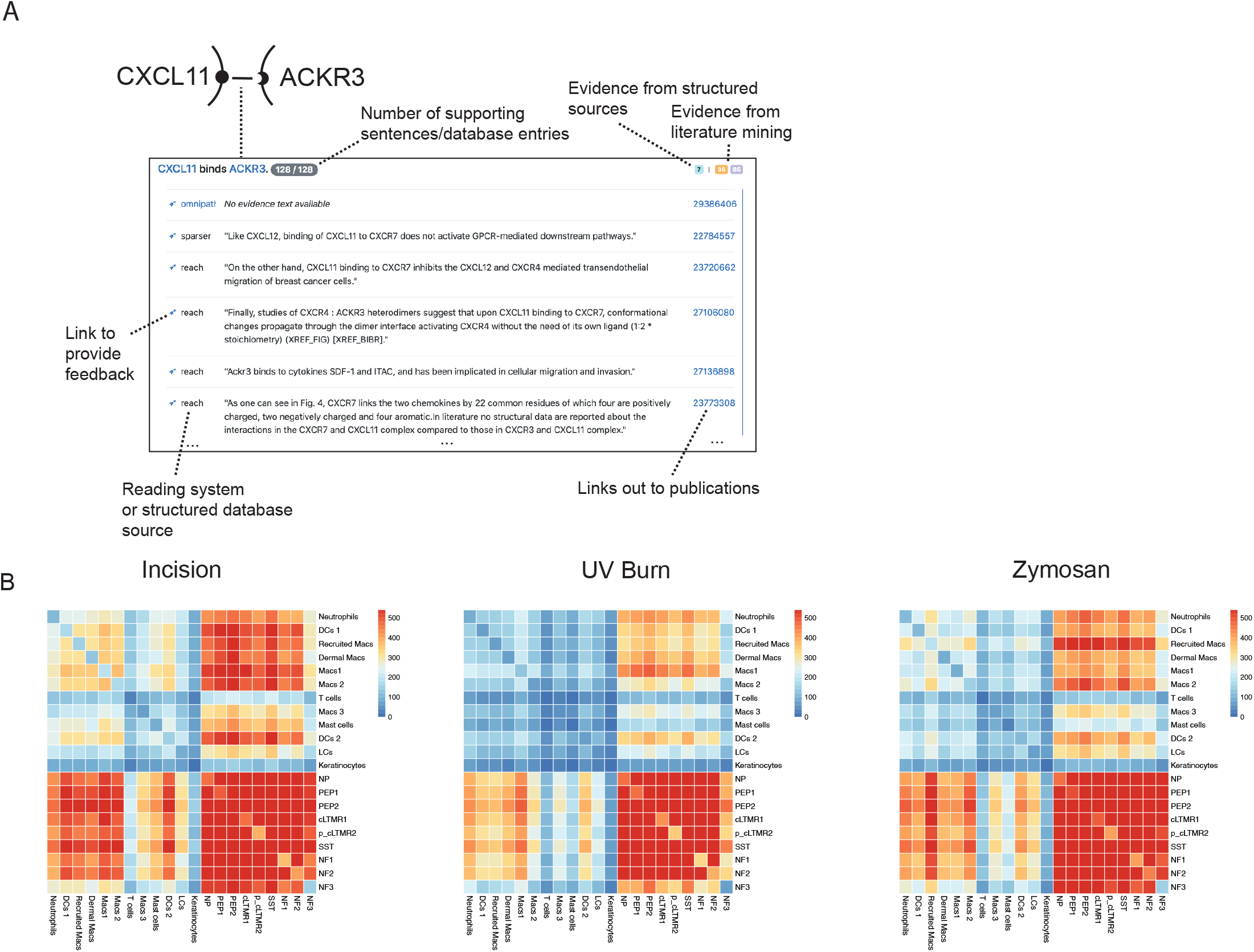
Constructing a receptor-ligand interactome using INDRA. (A) Sample of the web interface for browsing evidence behind the cell-cell interactome. Each interaction is displayed as a heading summarizing the interaction (“CXCL11 binds ACKR3”) with gene names linked to HGNC pages representing the gene. The total number of supporting pieces of evidence is shown as is the breakdown of this number by specific source: different structured databases or literature mining systems integrated with INDRA. Each row under the heading represents a distinct database entry or sentence from a publication, with each publication linked to its corresponding PubMed landing page. Each row also links to a curation page where feedback can be given on the correctness of the interaction. (B) Heatmap representing the number of significant interactions between given cell-cell pairs.

**Supplementary Figure 5.**
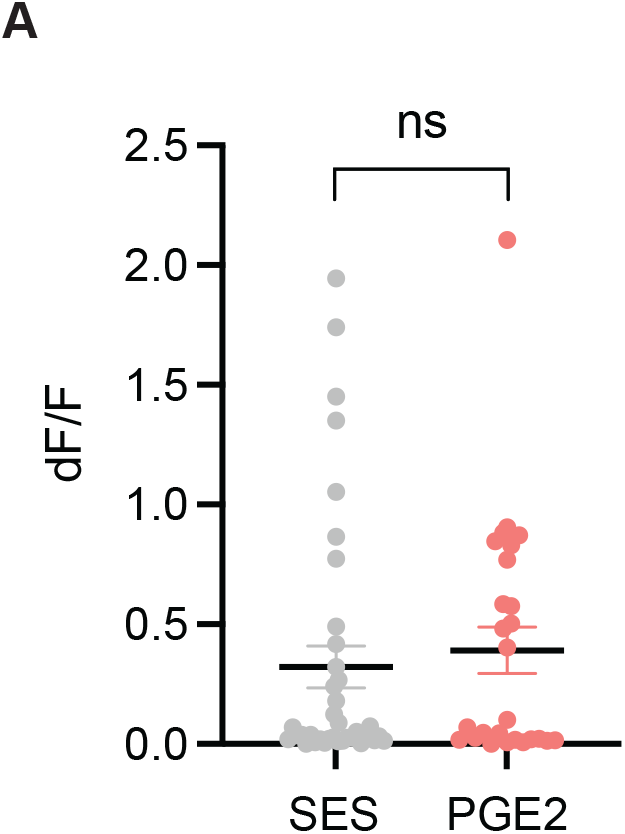
Cultured DRG neurons from female mice do not exhibit PGE2 sensitization of TRPV1. Fura-2-based calcium imaging was performed on cultured DRG neurons from healthy female mice. Treatment was performed the same as in Figure 6D. An unpaired t-test was performed. n.s. = not significant. Data is representative of two independent experiments.

## References

1. Basbaum, A.I., Bautista, D.M., Scherrer, G. & Julius, D. Cellular and molecular mechanisms of pain. Cell 139, 267–284 (2009).

2. Griffin, M.R. Epidemiology of nonsteroidal anti-inflammatory drug-associated gastrointestinal injury. Am J Med 104, 23S–29S; discussion 41S-42S (1998).

3. Becht, E. et al. Dimensionality reduction for visualizing single-cell data using UMAP. Nat Biotechnol (2018).

4. Kolter, J. et al. A Subset of Skin Macrophages Contributes to the Surveillance and Regeneration of Local Nerves. Immunity 50, 1482–1497 e1487 (2019).

5. Renthal, W. et al. Transcriptional Reprogramming of Distinct Peripheral Sensory Neuron Subtypes after Axonal Injury. Neuron 108, 128–144.e129 (2020).

6. Galli, S.J., Gaudenzio, N. & Tsai, M. Mast Cells in Inflammation and Disease: Recent Progress and Ongoing Concerns. Annu Rev Immunol 38, 49–77 (2020).

7. Kashem, S.W. et al. Nociceptive Sensory Fibers Drive Interleukin-23 Production from CD301b+ Dermal Dendritic Cells and Drive Protective Cutaneous Immunity. Immunity 43, 515–526 (2015).

8. Serhan, N. et al. House dust mites activate nociceptor-mast cell clusters to drive type 2 skin inflammation. Nat Immunol 20, 1435–1443 (2019).

9. Talbot, S. et al. Silencing Nociceptor Neurons Reduces Allergic Airway Inflammation. Neuron 87, 341–354 (2015).

10. Huang, J. et al. Circuit dissection of the role of somatostatin in itch and pain. Nat Neurosci 21, 707–716 (2018).

11. Honore, P. et al. Osteoprotegerin blocks bone cancer-induced skeletal destruction, skeletal pain and pain-related neurochemical reorganization of the spinal cord. Nat Med 6, 521–528 (2000).

12. Ley, K. & Morris, M. Signals for lymphocyte egress. Nat Immunol 6, 1215–1216 (2005).

13. Gyori, B.M. et al. From word models to executable models of signaling networks using automated assembly. Mol Syst Biol 13, 954 (2017).

14. John A. Bachman, B.M.G., Peter K. Sorger. Automated assembly of molecular mechanisms at scale from text mining and curated databases. Preprint at https://www.biorxiv.org/content/10.1101/2022.08.30.505688v1. (2022).

15. Efremova, M., Vento-Tormo, M., Teichmann, S.A. & Vento-Tormo, R. CellPhoneDB: inferring cell-cell communication from combined expression of multi-subunit ligandreceptor complexes. Nat Protoc 15, 1484–1506 (2020).

16. Turei, D., Korcsmaros, T. & Saez-Rodriguez, J. OmniPath: guidelines and gateway for literature-curated signaling pathway resources. Nat Methods 13, 966–967 (2016).

17. Turei, D. et al. Integrated intra- and intercellular signaling knowledge for multicellular omics analysis. Mol Syst Biol 17, e9923 (2021).

18. Bairoch, A. The ENZYME database in 2000. Nucleic Acids Res 28, 304–305 (2000).

19. Rodchenkov, I. et al. Pathway Commons 2019 Update: integration, analysis and exploration of pathway data. Nucleic Acids Res 48, D489–D497 (2020).

20. Adams, J.C. & Lawler, J. The thrombospondins. Cold Spring Harb Perspect Biol 3, a009712 (2011).

21. Isenberg, J.S. et al. CD47 is necessary for inhibition of nitric oxide-stimulated vascular cell responses by thrombospondin-1. J Biol Chem 281, 26069–26080 (2006).

22. Isenberg, J.S. et al. Thrombospondin-1 stimulates platelet aggregation by blocking the antithrombotic activity of nitric oxide/cGMP signaling. Blood 111, 613–623 (2008).

23. Aley, K.O. & Levine, J.D. Role of protein kinase A in the maintenance of inflammatory pain. J Neurosci 19, 2181–2186 (1999).

24. Taiwo, Y.O., Bjerknes, L.K., Goetzl, E.J. & Levine, J.D. Mediation of primary afferent peripheral hyperalgesia by the cAMP second messenger system. Neuroscience 32, 577–580 (1989).

25. Taiwo, Y.O. & Levine, J.D. Further confirmation of the role of adenyl cyclase and of cAMP-dependent protein kinase in primary afferent hyperalgesia. Neuroscience 44, 131–135 (1991).

26. Bhave, G. et al. cAMP-dependent protein kinase regulates desensitization of the capsaicin receptor (VR1) by direct phosphorylation. Neuron 35, 721–731 (2002).

27. Moriyama, T. et al. Sensitization of TRPV1 by EP1 and IP reveals peripheral nociceptive mechanism of prostaglandins. Mol Pain 1, 3 (2005).

28. Jain, A., Hakim, S. & Woolf, C.J. Unraveling the Plastic Peripheral Neuroimmune Interactome. J Immunol 204, 257–263 (2020).

29. Isenberg, J.S., Frazier, W.A. & Roberts, D.D. Thrombospondin-1: a physiological regulator of nitric oxide signaling. Cell Mol Life Sci 65, 728–742 (2008).

30. Christopherson, K.S. et al. Thrombospondins are astrocyte-secreted proteins that promote CNS synaptogenesis. Cell 120, 421–433 (2005).

31. Stein, C. & Lang, L.J. Peripheral mechanisms of opioid analgesia. Curr Opin Pharmacol 9, 3–8 (2009).

32. Klein, A.M. et al. Droplet barcoding for single-cell transcriptomics applied to embryonic stem cells. Cell 161, 1187–1201 (2015).

33. Satija, R., Farrell, J.A., Gennert, D., Schier, A.F. & Regev, A. Spatial reconstruction of singlecell gene expression data. Nat Biotechnol 33, 495–502 (2015).

